# Genome-scale metabolic model predicts mechanistic differences in metabolic gradient of spatially resolved small intestine villi

**DOI:** 10.1101/2023.06.22.546161

**Authors:** Joe Jongpyo Lim, Julia Yue Cui, Yuliang Wang

**Author notes:** **Address correspondence to:** Yuliang Wang, PhD, Research Assistant Professor, Institute for Stem Cell and Regenerative Medicine, University of Washington, Seattle, WA 98109, USA, Paul G. Allen School of Computer Science & Engineering, University of Washington, Seattle, WA 98195, USA, Joe Jongpyo Lim, PhD student, Department of Environmental and Occupational Health Sciences, University of Washington, Seattle, WA 98105, USA.

## Abstract

Studying the spatial metabolic gradient provides significant opportunities for understanding the spatial division of labor within the tissue microenvironment. Enterocytes have the capacity to perform serial oxidation and conjugation reactions for detoxification, making the small intestine important as one of the first-pass metabolic organs. Recently, the enterocyte metabolic gradient was found to exhibit differential metabolic preferences depending on its location in the villus. However, it remains unclear how metabolism mechanistically differs in enterocyte microenvironments.

To bridge this knowledge gap, we leveraged spatial transcriptomics data to (1) reconstruct genome-scale metabolic networks (GSMMs) that are location-specific, and (2) identify metabolic genes that may explain the differential susceptibility of different villus sections to diseases, diet, and other factors. We found that enterocytes at the bottom of the villus are enriched in genes related to intermediary metabolism, phase-I and -II metabolism, bile acid metabolism, and transporters. Comparing enterocyte GSMMs between the top and bottom of villi, we show that enterocytes at the top produce NAD and threonine more robustly compared to bottom enterocytes. Conversely, bottom enterocytes produce guanosine monophosphate (GMP) more readily than enterocytes at the top of the villus.

These metabolic differences may have implications for differential villi susceptibility to diseases such as neuroendocrine tumors, acute graft-versus-host disease, and nutritional perturbations such as high-fat diets. Taken together, our findings demonstrate in a mechanistic manner the metabolic differences of enterocytes in the small intestine, providing information that can be applied to additional disease states and inform therapeutics development.

**AUTHOR SUMMARY:** Significant metabolic gradients exist across different locations in a cell. Currently, bulk expression experiments, used in most animal studies and clinical trials, fail to describe this spatial variation due to the averaging effect in the tissue of interest. Recently, single-cell RNA sequencing technologies revealed heterogeneity within tissues. However, the underlying differences and mechanisms for the spatial metabolic variations remain understudied. In this research, we focus on the small intestine, which plays a critical role in nutrient absorption, distribution, and drug metabolism. The walls of the small intestine are composed of finger-like projections, called *villi*, that are composed of enterocytes. To investigate the potential mechanism underlying metabolic differences of enterocytes along the villi, we built and compared networks formed by gene-protein reaction relationships from spatially resolved bulk and scRNA-seq data. We found that enterocytes at the villus top more robustly produced key components in cell metabolism, such as nicotinamide adenine dinucleotide (nad) and threonine, whereas enterocytes at the villus bottom more robustly produced the nucleotide guanosine monophosphate (GMP). Our approach can be extended to the study of metabolic differences in other organs and diseases and to research the metabolism of specific compounds.

## INTRODUCTION

The small intestine is the major anatomical organ responsible for the enzymatic degradation of food and nutrient absorption. Once absorbed, nutrients pass from the enterocytes into enterohepatic circulation and enter the liver via the hepatic portal vein [1]. Although the small intestine is chiefly an absorptive organ, it can also metabolize endogenous and xenobiotic chemicals [2]. Similar to the liver, enterocytes have the capacity to undergo phase-I and -II metabolism, which are a series of oxidation and conjugation reactions for detoxification.

The relative location of cells in the same organ affects specialized metabolic capabilities due to their proximity and access to oxygen and cellular nutrients and differences in the cellular transcriptome. Thus, cells at different locations play differential roles in the organ. For example, in the liver, cytochrome P450s and glutathione S-transferase (GST) activities by hepatocytes occur near the periportal regions; glucose delivery, fatty acid oxidation, and sulfation reactions are favorably expressed around blood vessels [3]. Likewise, enterocytes have different, location-dependent metabolic capabilities, with genes involved in GST conjugation and acute phase response being enriched in the bottom sections of the villi and cell adhesion enriched in the top sections [4]. This cell microenvironment and metabolic landscape in the small intestine can give rise to differential susceptibilities from drug exposures or diseases.

The metabolic states of cells are increasingly recognized as playing critical roles in cell fate, homeostasis, and diseases [5–7]. *Genome-scale metabolic network models* (GSMMs) provide a systematic and mechanistic framework for modeling the connections among thousands of metabolites, enzymes, and transporters. GSMMs have been successfully used to model a wide variety of human diseases [8]. Small intestinal spatial heterogeneity and its different roles have been recently addressed using spatially reconstructed enterocytes via single-cell RNA sequencing (scRNA-seq); this research found that enterocytes at the lower end of the villus tended to have higher gene expression that encodes transporters related to amino acids, while enterocytic transporters at the higher end were more enriched in peptides, adipoproteins, and cholesterol [4]. However, it remains unclear how enterocyte microenvironment metabolism mechanistically differs and relates to diseases.

To address this knowledge gap of enterocyte spatial metabolic heterogeneity, we combined bulk and spatially resolved scRNA-seq reconstructed GSMMs for different sections of the small intestine. In particular, we developed a novel way to contrast the metabolic capabilities of different spatial locations (METHODS). Our goal was to leverage the powerful spatial transcriptomics data to identify metabolic genes that may explain the differential susceptibility of different villi sections to diseases and other factors.

## METHODS

### RNA sequencing (RNA-seq) processing and analysis

All data processing and analysis were conducted in the R environment [9]. RNA-seq counts data from laser-captured, micro-dissected mouse jejunum fragments were downloaded from Moor *et al*. [4]. Differential expression analysis was performed using DESeq2 [10]. Differentially expressed genes were defined as genes with adjusted *p*-values (Benjamini-Hochberg) < 0.1 in jejunum sections 2, 3, 4, and 5 compared to section 1. Differentially expressed genes with fold changes greater than absolute log_2_(1.5) were used as up-and-down-regulated genes. Relative to section 1, up-and-down-regulated genes were used as input for gene ontology enrichment using topGO [11], with all genes in the unfiltered expression table as the background.

Counts were converted to transcripts per million (TPM). A gene was considered as expressed if the expression sum exceeded the total number of samples. Then, genes with variance >1 across samples were kept. Principal component analysis was conducted on the filtered samples. Genes involved in xenobiotic metabolism and inflammation were categorized based on literature searches and extracted gene sets from the Gene Ontology consortium and the Kyoto Encyclopedia of Genes and Genomes (KEGG). Genes involved in intermediary metabolism were queried from work by Corcoran *et al*. [12]. Differentially expressed genes were matched with genes in each category of interest, namely inflammation, oxidative stress, intermediary metabolism, phase-I and -II metabolism, transporters, and bile acid metabolism. Proportions of differentially expressed genes in sections 1 and 5 were calculated and visualized as stacked bar plots.

### Metabolic reconstruction using updated mCADRE

Genes involved in critical metabolic processes (“metabolic genes” in this paper) were obtained by matching normalized and filtered genes with genes in draft metabolic reconstruction models for mice (iMM1415) [13] and its equivalent human model (Recon1) [14]. To investigate conserved genes and reactions, metabolic genes present in both models were used for further analysis.

Many algorithms, including mCADRE, aim to integrate transcriptomics data and metabolic networks. These metabolic models are developed to build one case-specific model at a time, independent of other models. However, a more common scenario is comparative model building. Oftentimes, to compare *in silico* metabolic differences, more than one metabolic network is built for comparison. In this case, a cutoff needs to be based on both absolute level and relative differences. The common threshold is 4 FPKM, which is the default in mCADRE; a gene leading to a reaction will be included in the model if the expression is 5 FPKM. However, in the differential comparison scenario, if the expression is 5 in section A and 30 in section B, then this reaction should still be considered non-core in section A even if it passed the absolute cutoff. For this reason, we developed a relative conversion method to contextualize and contrast different metabolic capabilities.

To compare the metabolic differences for sections 1 and 5, representing the bottom and top of the villus, respectively, we accounted for the absolute and relative gene expression to be included as core or non-core reactions in each metabolic model. The threshold was defined as 4 TPM, and, for each gene, the relative expression differences were calculated as

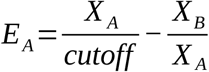

and

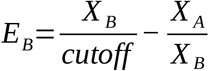

for

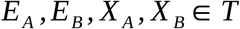

where T is a set of metabolic genes in sections A and B, and E_A_ and E_B_ are the relative expression differences for the gene of interest in model A and B, respectively. X_A_ is the expression of one gene in model A, and X_B_ is the expression of the same gene in model B, which is being compared to A. Expression values for all genes in both models (sections 1 and 5) were converted into absolute terms.

Using this relative thresholding, we defined a reaction as a core if the relative expression difference exceeded 4. Section-specific metabolic models were created using the mCADRE method [15], following the spatial metabolic network pipeline in Wang *et al*. [7]. Differential lethality analysis was conducted using the COBRA Toolbox [16] against sections 1 and 5 to predict key selective metabolic genes linked to reactions producing key metabolites for growth and survival. To maximize the robustness and decrease species-specific differences, all transcript isoforms for the same gene (e.g., x.1, x.2, x.3) in each model were knocked out when performing knockout iterations for differential lethal analysis. Upon knocking out each gene for all genes in each model, differentially lethal genes were defined as knocked-out genes that caused the growth rate ratio of section 1 (section 5) to be less than 0.2 and section 5 (section 1) to exceed 0.8. A secondary differential lethal analysis was conducted after deleting the primary differentially lethal genes in the model, where it was not lethal to predict alternative pathways in the non-lethal section to metabolize the key metabolite.

### Single-cell RNA-sequencing (scRNA-seq) data processing and analysis

Lgr5-eGFP -/-single-cell (sc) RNA-seq datasets were acquired from the NCBI GEO dataset browser (accessions GSM2644349 and GSM2644350 [17]. scRNA-seq data were read, filtered, and analyzed using Seurat 3.2.0 [18]. To increase reproducibility, cells were filtered, normalized, and processed following pipelines from Moor *et al*. [4]. Cell types such as goblet and neuronal cells were excluded, and only enterocytes and stem cell-like cells were included in the final analysis when comparing the gene expression of healthy cells.

The resulting cell clusters were further analyzed using the zonation reconstruction algorithm created by [4]. Briefly, landmark genes (from bulk RNA-seq) in each micro-dissected jejunum section were linked to the proportions of total cellular UMIs of each scRNA-seq cell cluster. Then, each cell cluster was assigned to one of 6 spatial villus zones based on the expression ranges of landmark genes and corresponding UMIs. A non-parametric permutation test was conducted to compute the *p*-values, which were adjusted using Storey’s method [19]. The resulting cell clusters were plotted using uniform manifold approximation and projection (UMAP) [20].

Alternative genes that are critical for a secondary pathway in synthesizing key metabolites in the non-lethal section were plotted along each section in the bulk RNA-seq and spatially inferred scRNA-seq data.

To evaluate the relevance of our model-predicted differentially lethal genes and alternative metabolic paths, RNA-seq, and microarray datasets were obtained from the following accession numbers: small intestine microarray data of mice on high-fat diets were downloaded from E-GEOD-29748; human small intestinal neuroendocrine primary tumor microarray data were obtained from E-GEOD-9576, and a mouse model of acute graft-versus-host disease small intestine RNA-seq data was obtained from E-MTAB-7765. Microarray intensity tables were read into R, log_2_-transformed, and quantile normalized. Differential expression was performed using limma [21]. The RNA-seq count table was read into R, and differential expression was conducted utilizing DESeq2 [10]. RNA-seq boxplot values were plotted as transcripts per million (TPM).

## RESULTS

### Spatially resolved genome-scale metabolic network models of the small intestine

Our study aimed to identify spatial metabolic heterogeneity in the small intestine, as shown in Fig. 1. We utilized the RNA-seq dataset from Moor *et al*. [4], which profiled the transcriptome of five laser-dissected sections of the small intestine (Fig. 1B). To understand the different metabolic landscapes and their potential mechanistic differences, we built GSMMs for sections 1 and 5, representing the top and bottom of the small intestinal villi. We performed systematic *in silico* single-gene knockout simulations and identified genes that, when knocked out, differentially affect the metabolic network states of sections 1 and 5 (Fig. 1A and METHODS). In each case, we also identified the secondary, alternative genes and metabolic paths that enable the section to bypass this primary gene in the section where it is not lethal (Fig. 1C).

**Figure 1.**
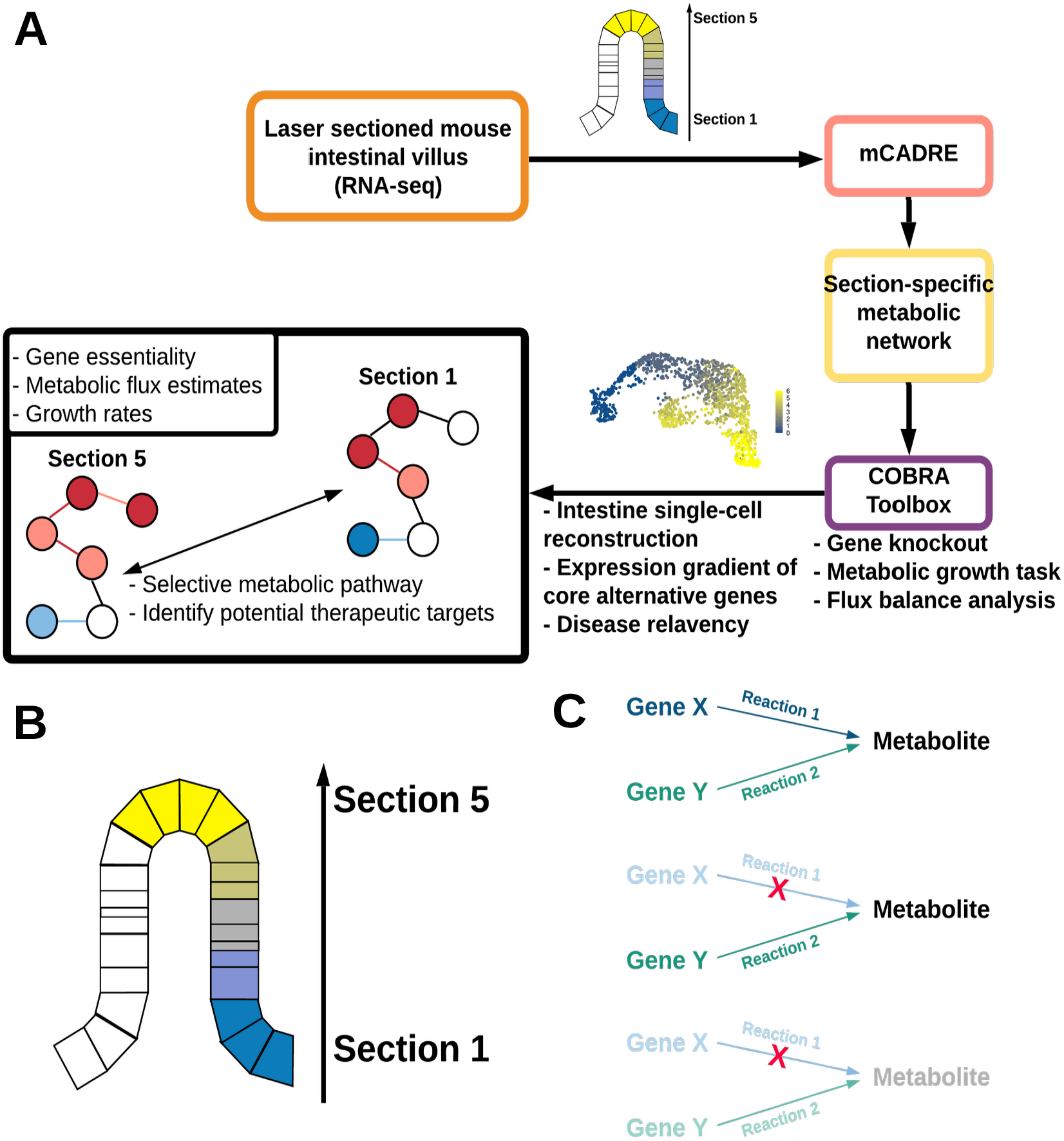
Study design, analysis workflow, and description of datasets. (A) A mouse villus was separated into five sections, and bulk RNA-seq was conducted. For sections 1 and 5, representing the two extreme locations of the villus, mCADRE was used to build section-specific metabolic models. *In silico* gene knockout, growth rate estimation, and flux balance analysis were performed using the COBRA Toolbox. Differentially lethal genes and alternative metabolic genes were found for each metabolic model and were cross-referenced with the spatially demultiplexed enterocyte single-cell RNA-seq dataset. Alternative metabolic genes’ disease relevance was then confirmed. (B) Laser-sectioned villi. The villi bottom, shown in blue, was denoted as section 1, and the villus top, shown in yellow, was defined as section 5, with sections 2, 3, and 4 in between. (C) Metabolic paths to metabolite production. Two metabolic paths produce a metabolite (top panel). If one gene (gene 1) linked to a core reaction (reaction 1) is down-regulated or knocked out, the reaction rate is not fast enough to produce the metabolite and would need to rely on gene 2/reaction 2 (middle panel). However, if gene 2 is not expressed in a particular cell, with the down-regulation of gene 1, the metabolite will not be produced in the cell (bottom panel). If the produced metabolites are necessary for cellular survival, the down-regulated reaction rate is referred to as *differentially lethal*.

### Spatial gradient in the small intestine

Using the small intestine scRNA-seq data from Yan *et al*. [17], we investigated the spatial metabolic gradient within this tissue. The villus zones were reconstructed by correlating the landmark expression genes of each section from the bulk RNA-seq data (described in [4]; see METHODS). The reconstructed villus zone contained a gradient of zero to six; zones 0 and 1 corresponded to the stem cell-like cells, and zones 2 to 6 were correlated with the enterocyte clusters (Fig. 2A and B). We performed principal component analysis (PCA) on the normalized and filtered bulk RNA-seq data. As shown in Figure 2C, a gene expression gradient was observed for the five villus sections; sections 1, 2, and 3 tended to cluster together, and sections 4 and 5 extended in the opposite direction.

**Figure 2.**
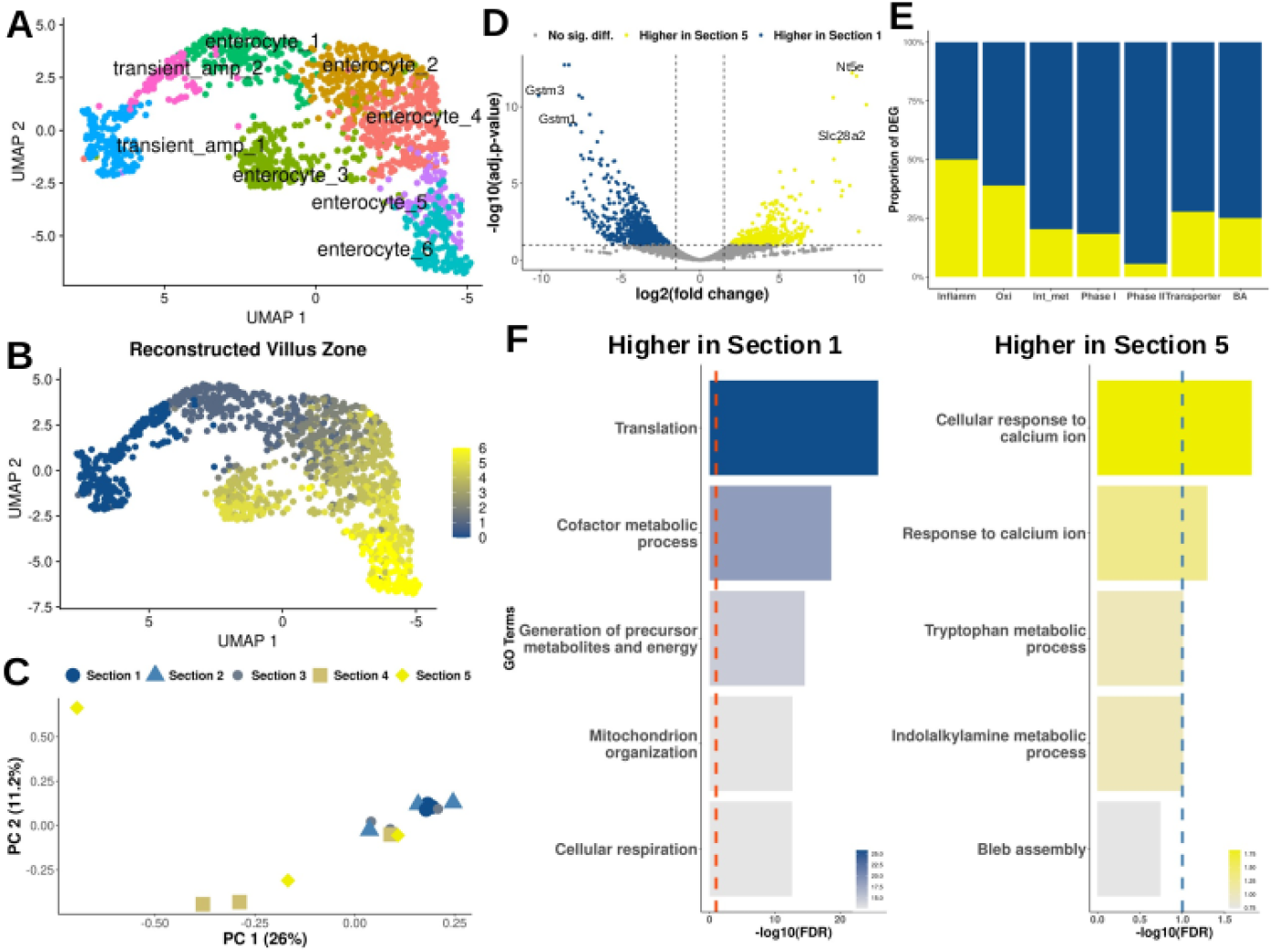
Metabolic differences between sections 1 and 5. (A) First two components of UMAP of the enterocyte single-cell RNA-seq dataset. (B) Spatially resolved representation of the enterocyte single-cell RNA-seq data, which correlates with the laser-dissected villi bulk RNA-seq dataset, shown in UMAP coordinates. (C) Principal component analysis on the five locations of the laser-sectioned villi. (D) Volcano plot of the total differentially expressed genes comparing sections 1 and 5. Blue shows genes expressed higher in section 1, while yellow indicates genes expressed higher in section 5. (E) Differentially expressed genes classified into metabolically relevant gene categories (from left to right, inflammation, oxidative stress, intermediary metabolism, phase I metabolism, phase II metabolism, transporters, and bile acid metabolism). The proportion, with respect to the total number of up-regulated genes, of sections 1 (blue) and 5 (yellow) were calculated and shown. (F) Gene ontology results showing the top 5 enrichment terms of section 1 (blue) and section 5 (yellow).

To understand the global metabolic gene expression differences between sections 1 and 5, we grouped differentially expressed genes into 7 major categories related to (1) enterocyte metabolism, and (2) investigated the proportion of genes in each category that were up-regulated in sections 1 or 5 compared to the other sections in each category (Fig. 2D). *Gstm1* and *3*, and *Nt5e* and *Slc28a2*, were among the top differentially expressed genes in sections 1 and 5, respectively (Fig. 2D and E, and Table S1). As shown in Figure 2A and Table S2, no major differences were found between sections 1 and 5 with respect to inflammation. However, section 1 contained a higher proportion of up-regulated genes in the oxidative stress (blue color), intermediary metabolism, phase-I and -II, transporters, and bile acid metabolism categories compared to section 5.

To group differentially expressed genes into biological processes in a more unbiased way, we performed gene ontology (GO) enrichment (Fig. 2F). Interestingly, we observed distinct roles between sections 1 and 5. Genes with higher expression in section 1 were enriched in translation, cofactor metabolism, precursor and energy metabolism, mitochondria organization, and cell respiration with high statistical significance. On the other hand, significantly enriched GO terms in section 5 included calcium ion response and tryptophan metabolism. Taken together, these results suggest that there are significant metabolic differences across the different locations of small intestinal villi.

Such metabolic differences may lead to differential susceptibility in dysregulated states, such as drug exposures or diseases. Therefore, to study the potential molecular mechanisms behind the metabolic differences between sections 1 and 5, we built section-specific GSMMs using a comparative approach (see METHODS). We identified ten genes that are essential for section 1 but not for section 5, and three genes that are essential for section 5 but not for section 1 (Table S3). We defined these genes as differentially lethal genes, focusing on three examples described below.

### Differential nucleotide metabolism

Nucleotide metabolism is one of the critical roles played by enterocytes. Small intestinal epithelial cells are among the first to encounter nucleotides from digestion [22]. These nucleotides are transported and metabolized by enterocytes, allowing further distribution to other cell types [22,23]. Guanosine monophosphate synthase (GMPS) is involved in converting ammonium and xanthosine 5’-phosphate into Guanosine monophosphate (GMP), which is a precursor in nucleotide synthesis. Enterocytes in both sections 1 and 5 rely on the GMPS enzyme for GMP synthesis (*Gmps* is not differentially expressed comparing sections 1 and 5, Fig 3A). Therefore, *in silico* knockout of GMPS resulted in selective lethality of enterocytes in section 1 due to the lack of an alternative pathway for synthesizing GMP. Interestingly, our model predicted that the knockout of GMPS was not lethal to enterocytes in section 5 because there is an alternative secondary pathway for GMP synthesis through 5’-nucleotidase Ecto (NT5E and also known as CD73, Fig. 3A). NT5E serves as an immune suppressor located on the cell surface and converts ATP, ADP, and AMP to adenosine [23] [24], [25]. NT5E can also convert GMP to guanosine (gsn) in the extracellular matrix (ECM). From the ECM to cytosol, GSNt4, a sodium symporting reaction, can import gsn, which can be converted to guanine (gua) via purine-nucleoside phosphorylase (PUNP3). GMP is then synthesized in the cytosol from gua via guanine phosphoribosyltransferase (GUAPRT).

**Figure 3.**
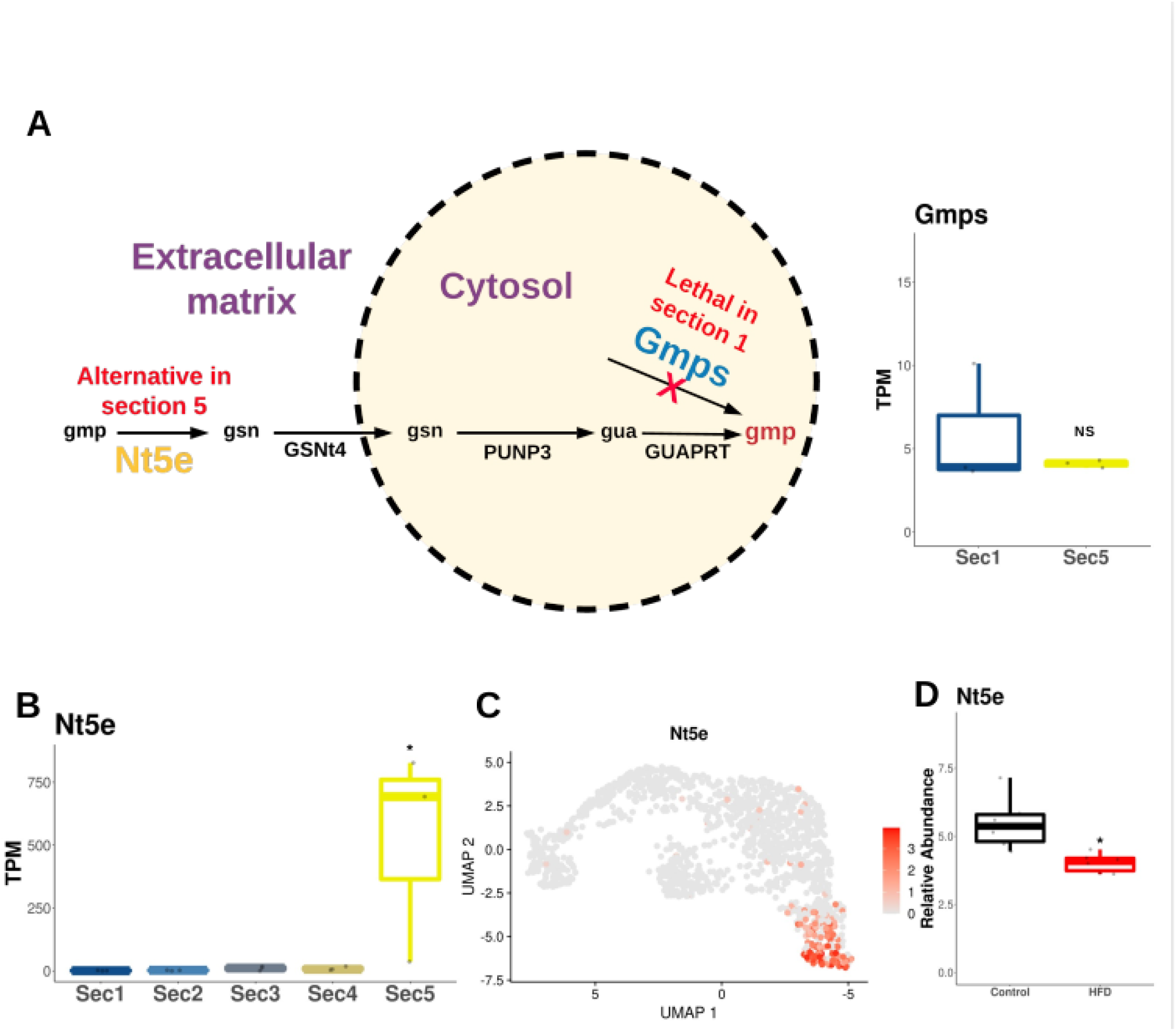
Mechanistic pathways of the lethal and alternative genes that produce guanosine monophosphate (gmp) and the expression patterns of the alternative gene in the bulk RNA-seq, single-cell RNA-seq, and mice fed with high-fat diets (HFDs). (A) Differentially lethal and alternative genes were predicted by comparing the metabolic models of sections 1 and 5 (see METHODS). Enterocytes in section 1 becomes lethal without *Gmps*, which is responsible for producing GMP in the cytosol. However, in section 5, *Nt5e* can produce gmp as an alternative pathway even when *Gmps* is knocked out in the model. (B) RNA expression gradient of *Nt5e* among the five laser-sectioned villi. (C) RNA expression patterns of *Nt5e* in the enterocyte single-cell RNA-seq data that is spatially correlated with the laser-sectioned villi bulk RNA-seq dataset. (D) Expression difference of *Nt5e* between mice fed with normal chow vs HFD in the small intestine. The asterisk shows significant differential expression.

The metabolic difference in GMP synthesis between two distinct sections of the villi is further supported by the mRNA expression levels of the key alternative pathway gene *Nt5e* in villi microsections, with higher average TPM levels in section 5 compared to section 1 (Fig. 3B). The selective expression pattern is also observed in spatially reconstructed enterocytes, with the inferred section 5 having a higher expression than section 1 (Fig. 3C). We also found that *Nt5e* expression is decreased in the small intestine of mice on an HFD (Fig. 3D). Combined with our model prediction, these findings suggest that an HFD will hinder the ability of section 5 to synthesize nucleotides in situations where GMPS activity is reduced because of its effect on reducing Nt5e expression.

### Differential NAD metabolism

Nicotinamide adenine dinucleotide (nad) is a necessary component in cells for universal metabolic processes, signaling, and post-transcriptional protein modifications [24]. *De novo* nad synthesis occurs through the enzyme glutamine-dependent nad synthetase, encoded by nad synthetase 1 (*Nadsyn1*), which was not differentially expressed comparing sections 1 and 5. *In silico* knockout of *Nadsyn1* is selectively lethal for section 5 but did not alter the growth rate in section 1. In section 1, a cluster of differentiation 38 (*Cd38*), involved in converting nad to cyclic-ADP-ribose (cADPR) [25], provides an alternative pathway to reuse existing nad from the ECM (Fig. 4A). Upon conversion of nad to nicotinamide (ncam) by Cd38, ncam is taken up by the NCAM uptake process (NCAMUP) to the cytosol. Ncam in the cytosol is transformed into nicotinamide mononucleotide (nmn) by NMN synthetase (NMNS). NMN can travel into the nucleus via nucleic pores (NMNtn) and is converted to nad via the enzyme nicotinamide-nucleotide adenylyltransferase (NMNATn). The resulting nad molecule can also enter the cytosol via gradient through pores.

**Figure 4.**
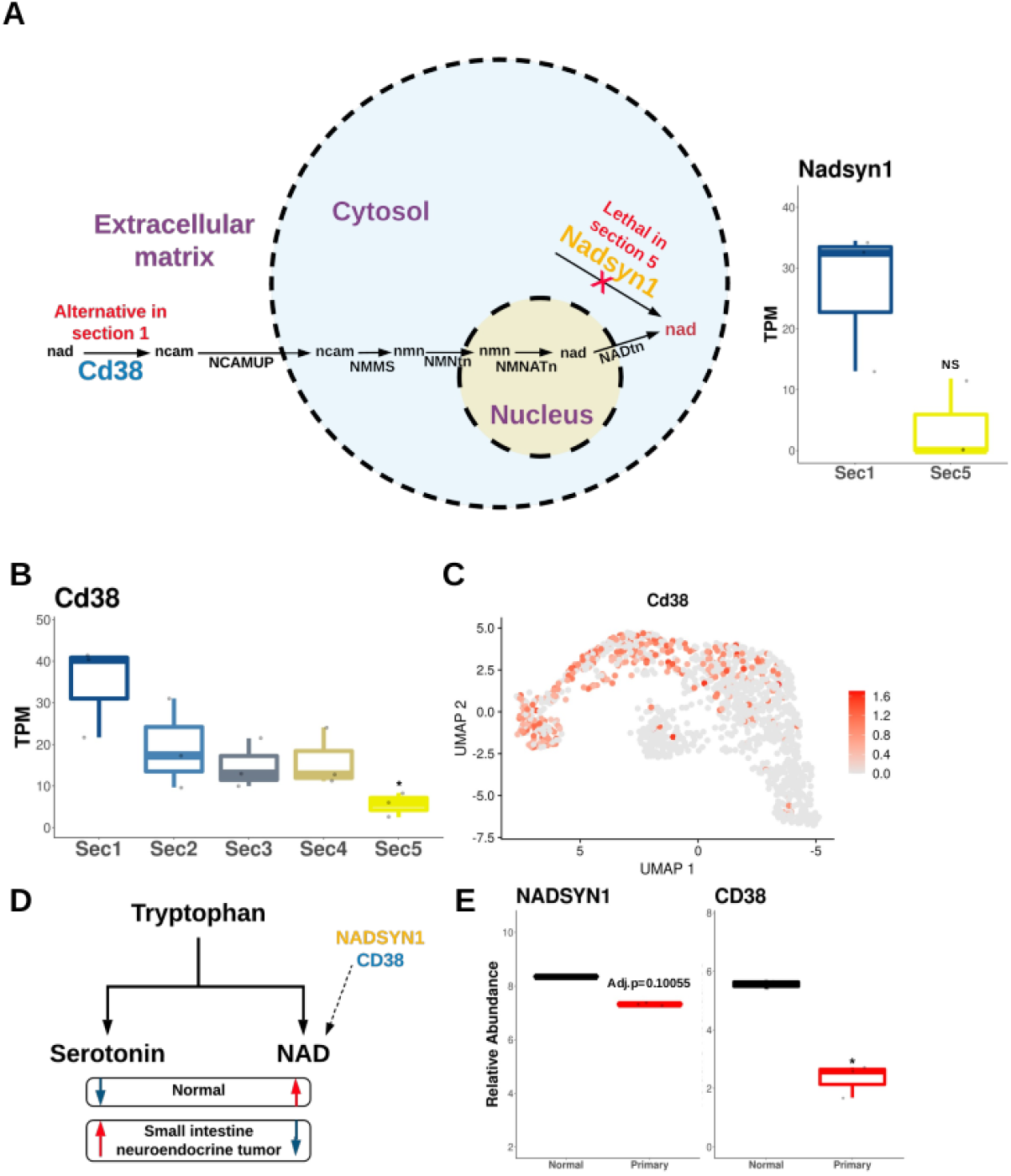
Mechanistic pathways of the lethal and alternative genes that produce nicotinamide adenine dinucleotide (nad); the expression patterns of the alternative gene in the bulk RNA-seq, single-cell RNA-seq; and human biopsy samples suffering from small intestine neuroendocrine primary tumors. (A) Differentially lethal and alternative genes were predicted by comparing the metabolic models of sections 1 and 5 (see METHODS). Enterocytes in section 5 become lethal without *Nadsyn1*, which is responsible for producing nad in the cytosol. However, in section 1, *Cd38* can produce nad as an alternative pathway even when *Nadsyn1* is knocked out in the model. (B) RNA expression gradient of *Cd38* among the five laser-sectioned villi. (C) RNA expression patterns of *Cd38* in the enterocyte single-cell RNA-seq data, which is spatially correlated with the laser-sectioned villi bulk RNA-seq dataset. (D) Serotonin and nad levels in normal vs small intestine neuroendocrine tumor patients. (E) Expression differences of *CD38* in human small intestine biopsy samples of normal (black) and primary small intestine neuroendocrine tumors (primary, red). The asterisk shows differential expression.

The expression of *Cd38* was the highest in section 1 and gradually decreased in section 5, which had the lowest expression from the laser-sectioned RNA-seq data (Fig. 4B). The expression gradient, as shown in the spatially decomposed single-cell data, shows that the expression level of *Cd38* is higher compared to section 5 (Fig. 4C).

Our prediction may be relevant in small intestine neuroendocrine tumors (NETs), i.e., slow-growing endocrine cell neoplasms that have become increasingly common over the past decades, that can accompany mood disorders [26,27]. It is known that altered tryptophan metabolism can lead to increased levels of serotonin and decreased levels of nad for patients with NETs compared to the normal condition (Fig 4D; adapted from Shah *et al*. [28]. Interestingly, the expression of *NADSYN1* in patients with small intestinal NETs decreased (adjusted *p*-value = 0.10055), suggesting that nad synthesis is lower for NET patients. Notably, *CD38*, the core gene for the alternative pathway in nad metabolism, was down-regulated for NET patients, which may indicate that lower nad levels in patients with NET are due to the dysregulated and differential metabolism of enterocytes in section 1 (Fig. 4E).

### Differential amino acid transport across the small intestine

Our global analysis revealed that many transporters are enriched in section 1 (Fig. 2A). Interestingly, our model predicted that solute carrier family 36 member 1 (*Slc36a1*), which encodes a protein that transports small amino acids, is essential for section 5 but not 1. *Slc36a1* can transport glycine into the cytosol from the ECM, and threonine can be imported inside the cytosol via the threonine-glycine exchange reaction (THRGLYexR). The villus section 5 model is predicted to rely on threonine transport by glycine exchange and lacks other threonine transport pathways. On the other hand, section 1 is not sensitive to *Slc36a1* knockout: in addition to glycine-threonine symport, solute carrier family 43 member 2 (*Slc43a2*) is predicted to be the key amino acid transporter for the alternative threonine transport pathway in section 1. Cysteine and threonine can be transported between the cytosol and the ECM via the cysteine-threonine sodium-dependent exchange reaction (THRCYSNAexR), resulting in cytosolic threonine. Given the deletion of Slc36a1, the transport of cysteine into enterocytes in section 1 by *Slc43a2* can drive THRCYSNAexR, thereby supplying threonine for enterocytes in this section.

Interestingly, in the bulk RNA-seq dataset, the expression of *Slc43a2* was the lowest in section 5 relative to the other four sections, all of which had approximately 1.5-fold higher expression levels of the gene (Fig. 5B). The plateau effect was also observed in the single-cell data, for which the expression of Slc43a2 was significantly higher in all sections but 5 (Fig. 5C).

**Figure 5.**
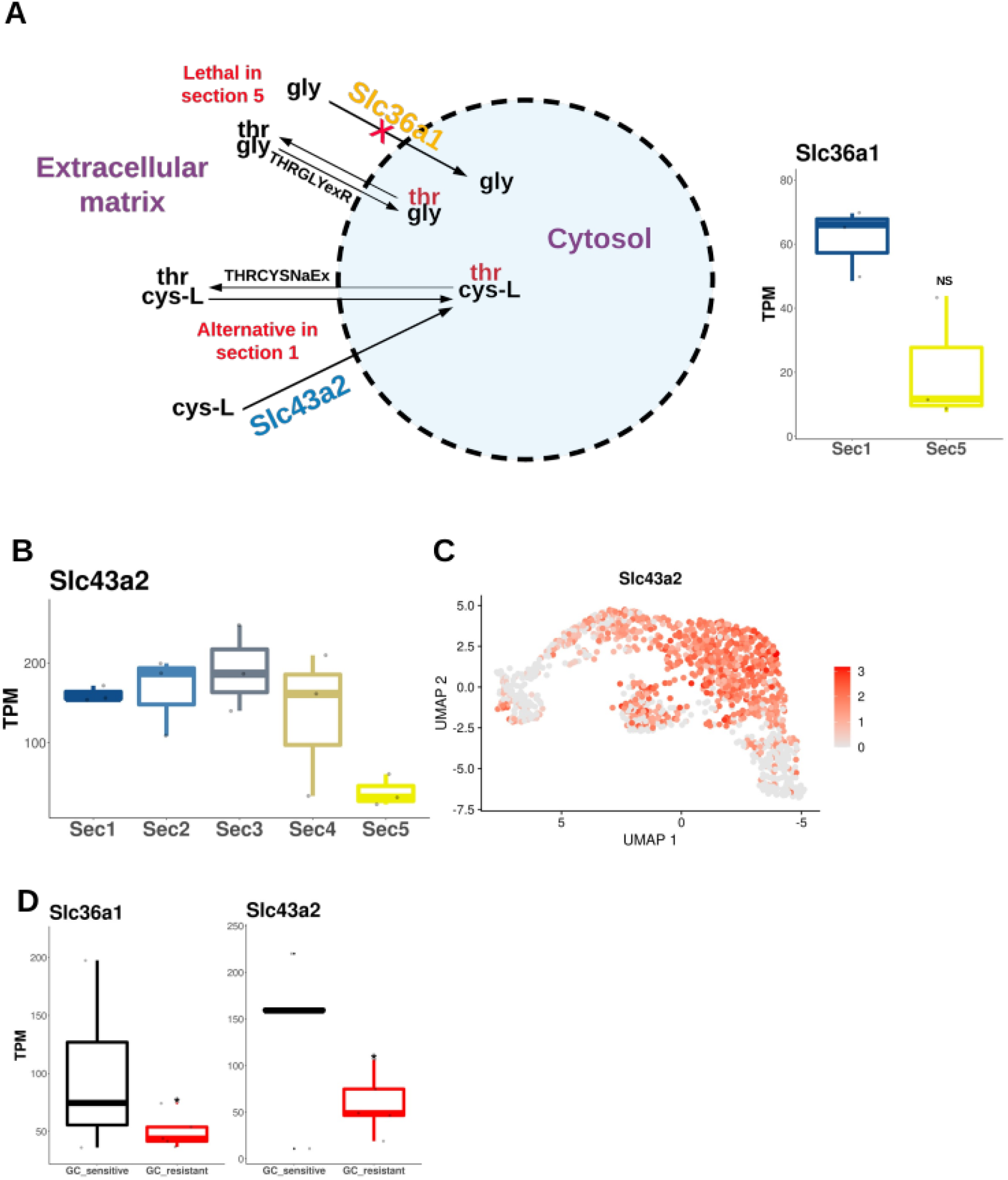
Mechanistic pathways of the lethal and alternative genes that produce threonine (thr), and the alternative gene expression patterns in the: bulk RNA-seq, single-cell RNA-seq, and acute graft-vs-host disease-induced mice. (A) Differentially lethal and alternative genes were predicted by comparing the metabolic models of sections 1 and 5 (see METHODS). Enterocytes in section 5 become lethal without *Slc36a1*, which is responsible for transporting thr into the cytosol. However, in section 1, *Slc43a2* can transport L-cysteine to produce threonine as an alternative pathway even when *Slc36a1* is knocked out in the model. (B) The RNA expression gradient of *Slc43a2* among the five laser-sectioned villi. (C) RNA expression patterns of *Slc43a2* in the enterocyte single-cell RNA-seq data, which is spatially correlated with the laser-sectioned villi bulk RNA-seq dataset. (D) Expression differences of *Slc43a2* in mice induced with acute graft-vs-host disease (red) compared with the control (black). The asterisk shows differential expression.

Our model prediction may be relevant in graft-versus-host disease (GvHD). GvHD is an inflammation disorder in immune cells (e.g., in bone marrow, stem cells, tissues, and solid organs) upon transplantation due to an innate immune response from detecting the graft as foreign [31]. It can lead to cytokine production and target tissue damage, which can result in health problems in the recipient. Strikingly, in the small intestine of mice induced with acute GvHD (aGvHD), both the primary gene for threonine transport, *Slc36a1*, and the key alternative gene, *Slc43a2*, were down-regulated in glucocorticoid (GC) resistant mice; this may indicate that the disease preferentially targeted section 1 enterocytes via threonine metabolism (Fig. 5D). In combination with model prediction, these results suggest that aggravated aGvHD due to GC resistance can lead to altered threonine metabolism due to the selectivity of enterocytes following metabolic differences.

## DISCUSSION

The spatial location of cells can profoundly affect their function. Studying the spatial gradient of cells within an organ can help us understand the cell microenvironment and its changes in complex diseases and therapeutics. For example, the tumor microenvironment contains many different cell types with different nutrient needs and metabolisms [29–31]. The small intestine, in addition to its nutrient-absorptive property, is critical for metabolizing endogenous compounds and xenobiotics that involve phase-I and -II metabolic pathways [2]. In addition, the small intestine is closely connected to other metabolically important organs, such as the liver. Recently, the expression gradient in small intestinal villi was reported [4]. *By comparing the two ends of small intestinal villi, we identified for the first time their potential metabolic differences at the mechanistic level*. We established that selective metabolic difference paths exist, both globally and mechanistically, depending on the location in the villus (Figs. 1-5).

Our global analysis shows that enterocytes at villi’s bottom, compared to villi’s top, are more enriched genes involved in intermediary, phase-I and -II, and bile acid metabolism as well as in transporters (Fig. 2A). In addition, we show that GO enrichment terms are related to cofactor precursor metabolism in section 1, whereas GO terms are involved in calcium response in section 5 (Fig. 2C). Such difference may be due to the selective expression of transcription factors important in endogenous molecules and xenobiotic biotransformation, such as *Esrra* (energy production), *Hnf4g* (steroid metabolism), *Nr1i3* (xenobiotic metabolism), *Vdr* (vitamin D metabolism), and *Nfe2l2* (oxidative stress) (Table S3). By comparing the GSMMs between small intestinal villi sections 1 and 5, we show that enterocytes at the top of villi are enriched with alternative pathways to produce nad and thr inside the cytosol compared to enterocytes at the bottom (Fig. 6), with specific examples of the changes from human NET and aGvHD in a mouse model (Fig. 4 and 5). Conversely, supported by mice fed with HFD, enterocytes at the bottom can produce GMP more readily than those at the villi’s top (Fig. 3). Taken together, our findings establish the presence of metabolic differences of enterocytes in the small intestine in a mechanistic manner, which may prove relevant to additional disease states and provide information to aid in the development of therapeutics (Fig. 6).

**Figure 6.**
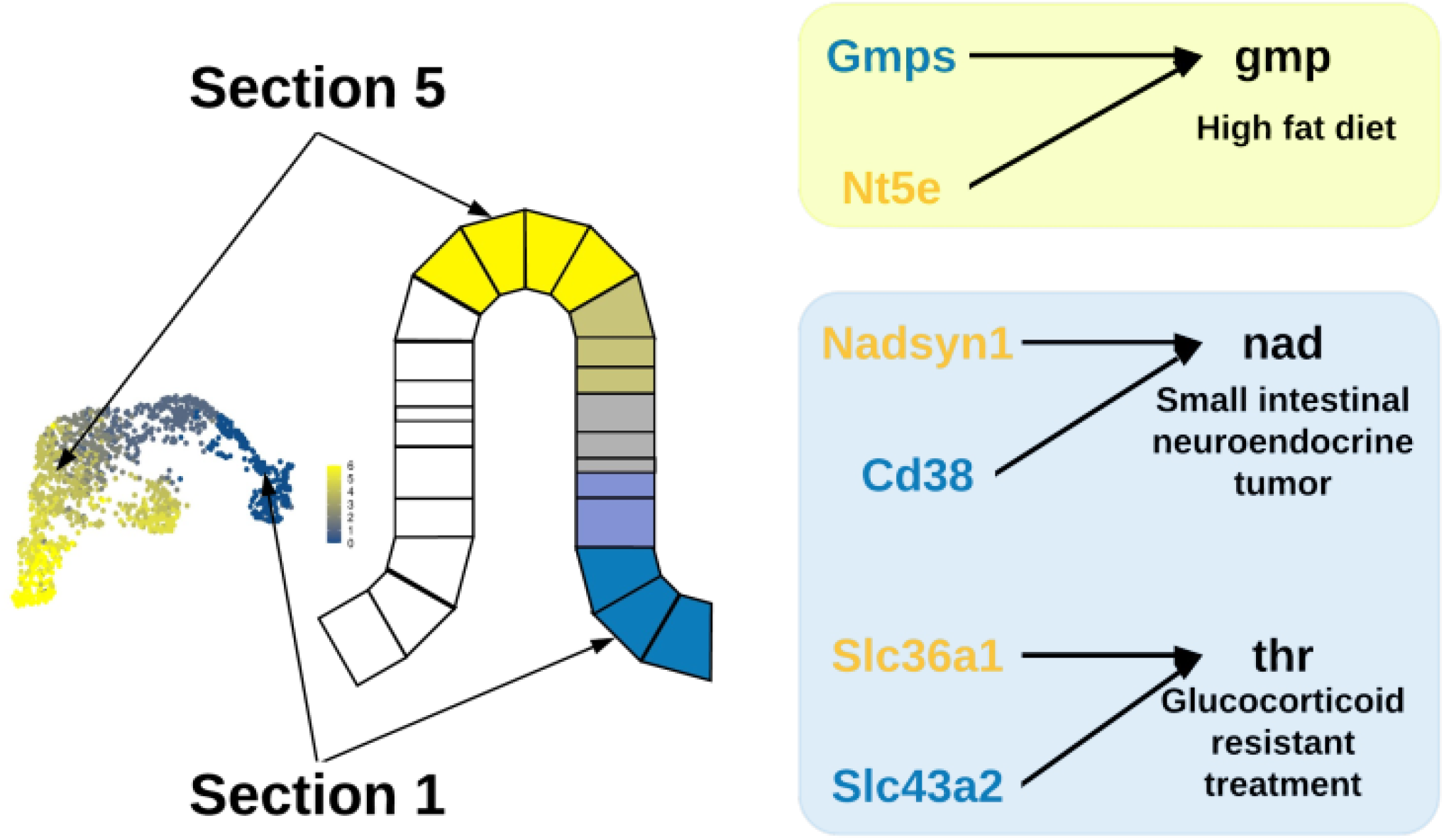
Summary figure of study findings. Differentially lethal genes were predicted by comparing the growth rate ratios of metabolic models of sections 1 and 5. Alternative genes were estimated by performing a second-degree *in silico* knockout of metabolic genes related to the produced metabolite by the differentially lethal genes. (Left) A bulk RNA-seq of laser-sectioned intestinal villus and spatially reconstructed single-cell RNA-seq villus zones. (Right) Lethal and alternative genes for each intestinal section are represented in unique colors (blue – section 1, yellow – section 5). From our results, we predicted *Gmps* is differentially lethal in section 1, and *Nadsyn1* and *Slc36a1* are differentially lethal in section 5. We found that *Nt5e, Cd38*, and *Slc43a2* are the key metabolic alternative genes to produce gmp, nad, and thr, respectively. The altered metabolites from *in silico* knockout experiments were further evidenced by disease phenotypes from high-fat diets, small intestinal neuroendocrine primary tumors, and glucocorticoid-resistant treatment from acute graft-vs.-host disease.

The generated models focused on enterocytes, excluding other cell types in the small intestine. Although scRNA-seq greatly improves the resolution within an organ by measuring cell-type-specific gene expression, the resulting expression matrix is sparse. These limitations warrant future work focusing on transforming sparse expression matrices to build cell-type-specific metabolic models. Furthermore, our findings lack direct experimental support, and the metabolic models were built without the integration of metabolite or protein information; these factors increase the potential for missing additional metabolic differences in the model. However, we recapitulated the metabolic gradient and differences between the two villi sections using gene expression and support our findings in additional disease states (Fig. 3-5). Furthermore, our study used a powerful approach, which features the highest sequencing depth per sample, to link bulk RNA-seq and spatially resolved scRNA-seq data of enterocytes, showing potential mechanistic differences in the metabolic gradient along small intestinal villi.

The gut microbiome, i.e. microorganisms that live inside the small and large intestines, has been increasingly recognized as contributing to host metabolism and is known to be dysregulated by disease and toxicant exposures [31–33], which its metabolic capabilities have been previously modeled [34,35]. Future work should connect our understanding of the metabolic gradient of enterocytes with other metabolically important organs, such as the gut microbiome and the liver. Our novel modeling approach, which considers the relative difference of metabolic genes, can be applied to a variety of fields, such as the cancer microenvironment and differential susceptibility to drugs across cell types and locations. This holds promise for investigating metabolic differences in a mechanistic manner, thus revealing the potential mechanism of condition-specific metabolic alterations.

